# Heterotopic expression of respiratory-associated genes suggests a developmental basis for pleopodal gill evolution in isopod crustaceans

**DOI:** 10.64898/2026.11.09.765247

**Authors:** Naoto Inui, Akifumi Yao, Kenji Fukushima, Toru Miura

## Abstract

**Background:** The evolution of organs in new anatomical positions contributes to morphological diversity and offers opportunities to examine developmental evolution. Crustacean gills provide a useful comparison: isopods bear pleopodal gills on abdominal appendages, whereas many other crustaceans possess thoracic epipodal gills. Here, we combined comparative anatomy and multiplex *in situ* hybridization chain reaction to examine embryonic expression of the respiratory-associated genes *trachealess* (*trh*) and *ventral veins lacking* (*vvl*) in the aquatic isopod *Gnorimosphaeroma* cf. *hoestlandti* and the terrestrial isopod *Porcellio scaber*. We also examined the posterior Hox genes *abdominal-A* and *Abdominal-B* (*AbdB*) to assess regional pleonal patterning.

**Results:** In both species, *trh* and *vvl* expression occurred in developing pleopodal regions corresponding to gill-bearing tissues. In *G.* cf. *hoestlandti*, prominent expression extended into both endopodal and exopodal regions of pleopods 4–5, whereas in *P. scaber*, it was concentrated in the endopods of pleopods 3–5. In both species, *trh* showed more restricted posterior pleopodal expression than *vvl*, and posterior expression domains overlapped spatially with *AbdB*-expressing regions.

**Conclusions:** Compared with thoracic epipodal expression reported in other crustaceans, these patterns differ at two anatomical levels: along the body axis (thorax versus pleon) and among appendage structures (epipods versus endopods/exopods). These differences suggest that the acquisition of heterotopic *trh* and *vvl* expression in isopod pleopods may have contributed to the evolution of pleopodal gills. Moreover, the different expression patterns in the two isopods suggest subsequent evolutionary changes in the spatial distribution of these genes, potentially associated with the diversification of pleopodal gills.

## Introduction

Morphological evolution can involve changes not only in organ structure but also in where organs develop. Arthropods provide a particularly clear framework for studying such positional diversification because their serially repeated appendages have diversified extensively in form and function along the body axis (Bruce, 2021; Jockusch, 2017). Crustacean gills exemplify this positional diversity. In many malacostracans, gills are associated with thoracic appendages, including epipods and tissues around the appendage base. Even among epipod-associated respiratory systems, the segmental distribution of gills varies among crustacean lineages. For example, lophogastrids bear branchial epipodites on multiple thoracopods, whereas in cumaceans and tanaidaceans epipods are restricted to the maxilliped, where they contribute to respiration and ventilation (Boxshall & Jaume, 2009; Tan & Monteiro, 2025; Wirkner & Richter, 2013). Such diversity can occur at multiple anatomical scales, from differences among serial appendages to differences in the structures that bear respiratory tissue. Understanding the developmental basis of this variation is therefore important for explaining the evolution of gill organization.

The transcription factors *trachealess* (*trh*) and *ventral veins lacking* (*vvl*; also known as *drifter* or *ventral veinless*) offer a molecular entry point into this question. In the model insect *Drosophila melanogaster*, Trh is essential for tracheal development, whereas Vvl contributes to tracheal cell differentiation and migration (Anderson et al., 1995; Chung et al., 2011; Isaac & Andrew, 1996). Orthologues of these genes are also expressed in respiratory or osmoregulatory structures in crustaceans. In *Artemia* branchiopods, *trh* is expressed in the larval salt gland and thoracic epipod, and *vvl* is also expressed in the epipod (Mitchell & Crews, 2002; Wang et al., 2012); in the amphipod *Parhyale hawaiensis*, both genes are expressed in developing thoracic epipodal gills (Franch-Marro et al., 2006). Together, these observations suggest that *trh* and *vvl* are deployed across anatomically distinct respiratory and osmoregulatory structures.

Isopods provide an informative system in which to examine this possibility because their gills occur on abdominal appendages (pleopods) rather than thoracic epipods (Fig. 1A). Their pleopods consist of inner endopods and outer exopods, one or both of which can contribute to respiration and osmoregulation. Pleopodal respiratory organization has diversified further in terrestrial isopods, in which gills coexist with lung-like air-breathing structures (Inui & Miura, 2024; Tan & Monteiro, 2025). Previous studies further suggest that *trh* and *vvl* are associated with pleopodal gills: embryonic *vvl* expression has been reported in posterior pleopods, and adult transcriptomic analyses detected higher expression of both *trh* and *vvl* in endopods than in exopods (Abzhanov & Kaufman, 2000b; Inui et al., 2025). However, the embryonic expression of *trh* has not been investigated or compared between isopods with different gill arrangements. Such comparisons can reveal how variation in the spatial deployment of these genes relates to evolutionary variation in gill position.

**Figure 1.**
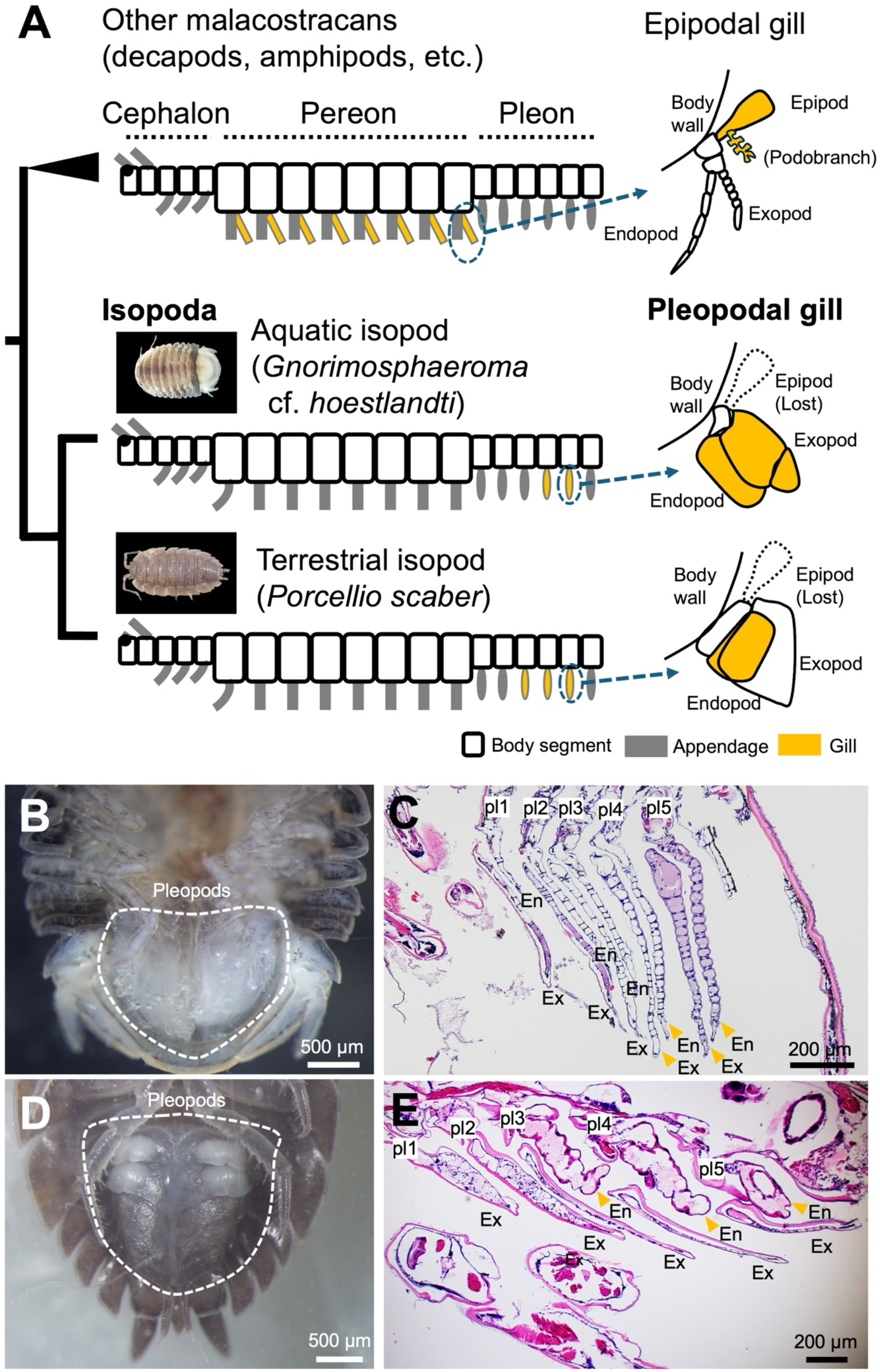
Pleopodal gills in isopod crustaceans. (A) Schematic comparison of appendage-associated gills in malacostracans and the two focal isopods. Epipodal gills are illustrated on thoracic appendages, whereas the focal isopod gills occur on pleopodal rami. Yellow indicates gill regions. The branching diagram summarizes the selected anatomical comparisons. (B, C) Ventral view of the adult pleon and histological section through the pleopods of *Gnorimosphaeroma* cf. *hoestlandti*. (D, E) Corresponding views of *Porcellio scaber*. White dashed outlines in B and D delimit the pleopodal area. Orange arrowheads in C and E indicate the posterior gill regions highlighted for comparison. En, endopod; Ex, exopod; pl1–pl5, pleopods 1–5. Scale bars: 500 µm in B and D; 200 µm in C and E. Schematics are not to scale.

Here, we examined the embryonic expression of *trh* and *vvl* in the aquatic isopod *Gnorimosphaeroma* cf. *hoestlandti* and the terrestrial isopod *P. scaber* using multiplex *in situ* hybridization chain reaction (HCR). The two species exhibit different pleopodal gill arrangements: in *G.* cf. *hoestlandti*, gill-bearing regions occur in both the endopods and exopods of the posterior pleopods, whereas in *P. scaber*, the posterior gills are confined to the endopods (Fig. 1). We characterized these gill-bearing tissues anatomically to provide a reference for interpreting the expression domains. To gain insight into the developmental regulatory genes associated with the pleonal expression of *trh* and *vvl*, we also examined the posterior Hox genes *abdominal-A* (*abdA*) and *Abdominal-B* (*AbdB*), given the roles of Hox genes in axial patterning in crustaceans (Brena et al., 2005; Martin et al., 2016). By relating *trh* and *vvl* expression to distinct gill-bearing rami and comparing these patterns with published thoracic epipodal expression in other crustaceans, we explored the possibility that changes in the spatial deployment of respiratory-associated genes contributed to the evolution of pleopodal gills.

## Results

### Pleopodal gill differentiation in the aquatic and terrestrial isopods

To establish an anatomical reference for the gene-expression analyses, we first examined pleopodal tissues in the two focal isopods. Both species possessed five pairs of pleopods on the ventral side of the pleon (Fig. 1B–E). In *Gnorimosphaeroma* cf. *hoestlandti*, all five pairs consisted of flattened endopods and exopods. Histological sections revealed conspicuous differentiation of the posterior pleopods: pleopods 4 and 5 had more extensively folded epithelia and thinner cuticles than the anterior pleopods (Fig. 1C, orange arrowheads). In *Porcellio scaber*, the posterior three pairs of pleopods were differentiated into exopods and endopods. The exopods were flattened and covered by a relatively thick cuticle, whereas the endopods consisted of thicker tissue covered by a thin cuticle (Fig. 1E, orange arrowheads).

These observations identified anatomically distinct posterior pleopodal regions in the two species: differentiated endopodal and exopodal regions in *G.* cf. *hoestlandti* and posterior endopodal regions in *P. scaber*. We used these regions as anatomical references for localizing embryonic gene expression.

### Embryonic development of the aquatic and terrestrial isopods

Embryonic development in *G.* cf. *hoestlandti* and *P. scaber* showed broadly comparable morphological features, allowing the 20-stage developmental staging system of Wolff (2009) to be applied to both species (Fig. S1). For the gene-expression analyses, we examined embryos at stages 11–18, grouping them into early (stages 11–13), middle (stages 14–16), and late (stages 17–18) intervals within the examined developmental period. During these stages, pleopods first appeared as limb buds with two distal outgrowths, which subsequently elongated and differentiated into distinct exopods and endopods (Figs. 2–5).

**Figure 2.**
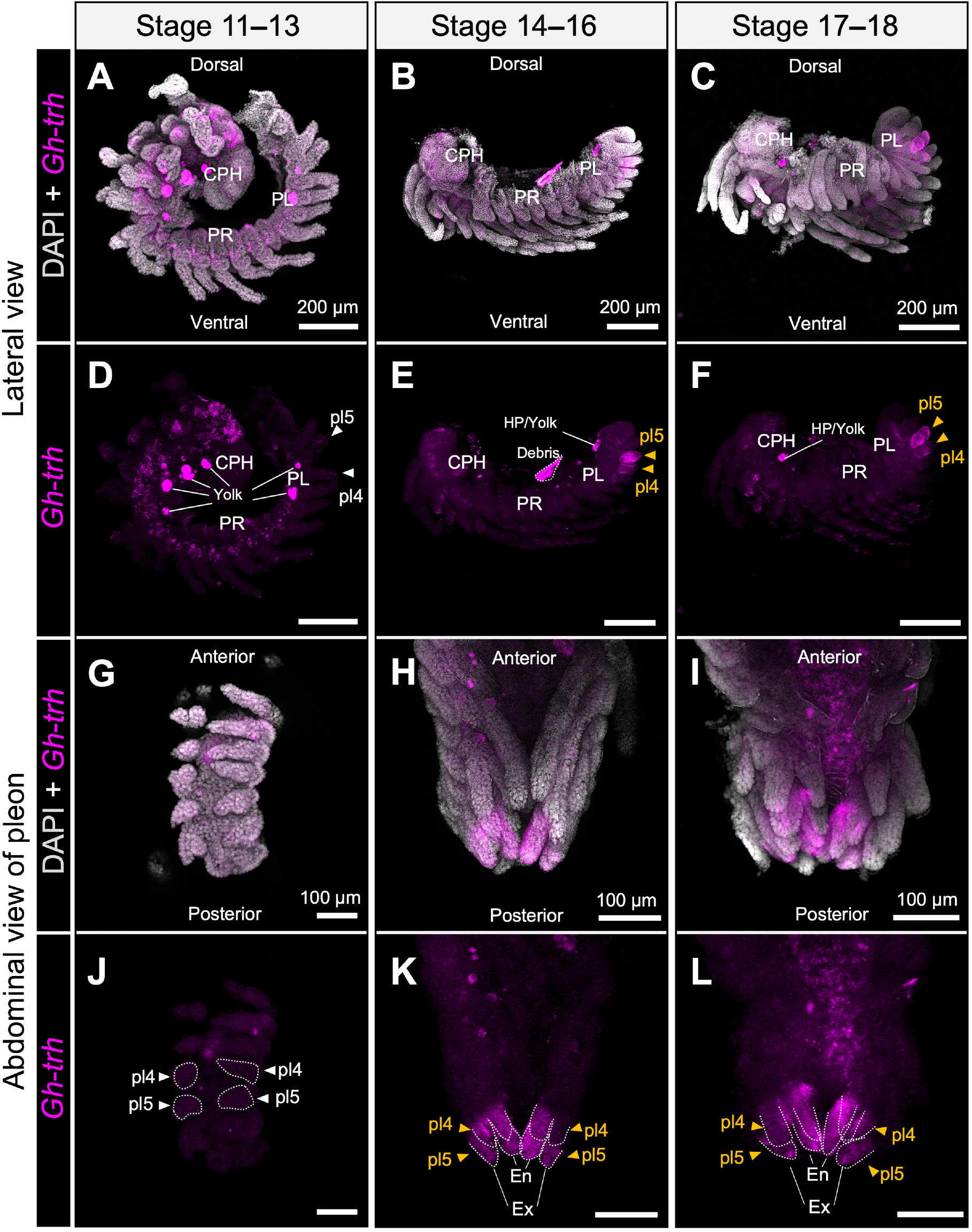
Embryonic expression of *trachealess* in *Gnorimosphaeroma* cf. *hoestlandti*. HCR localization of *Gh-trh* across stages 11–13 (A, D, G, J), 14–16 (B, E, H, K) and 17–18 (C, F, I, L). (A–F) Lateral views, with DAPI and *Gh-trh* merged in A–C and the *Gh-trh* channel shown in D–F. (G–L) Abdominal views of the pleon, with merged images in G–I and the *Gh-trh* channel in J–L. *Gh-trh* signal is magenta and DAPI is gray. Arrowheads identify pleopods 4 and 5; white arrowheads mark their positions in the early specimens, and orange arrowheads highlight the more clearly localized later domains. Dotted outlines in J–L delimit selected pleopodal regions. Structures labeled yolk, HP/yolk or debris are excluded from expression interpretation. CPH, cephalon; PR, pereon; PL, pleon; En, endopod; Ex, exopod; pl, pleopod; HP, hepatopancreas. Orientation is indicated in the panels. Scale bars: 200 µm in A–F; 100 µm in G–L.

**Figure 3.**
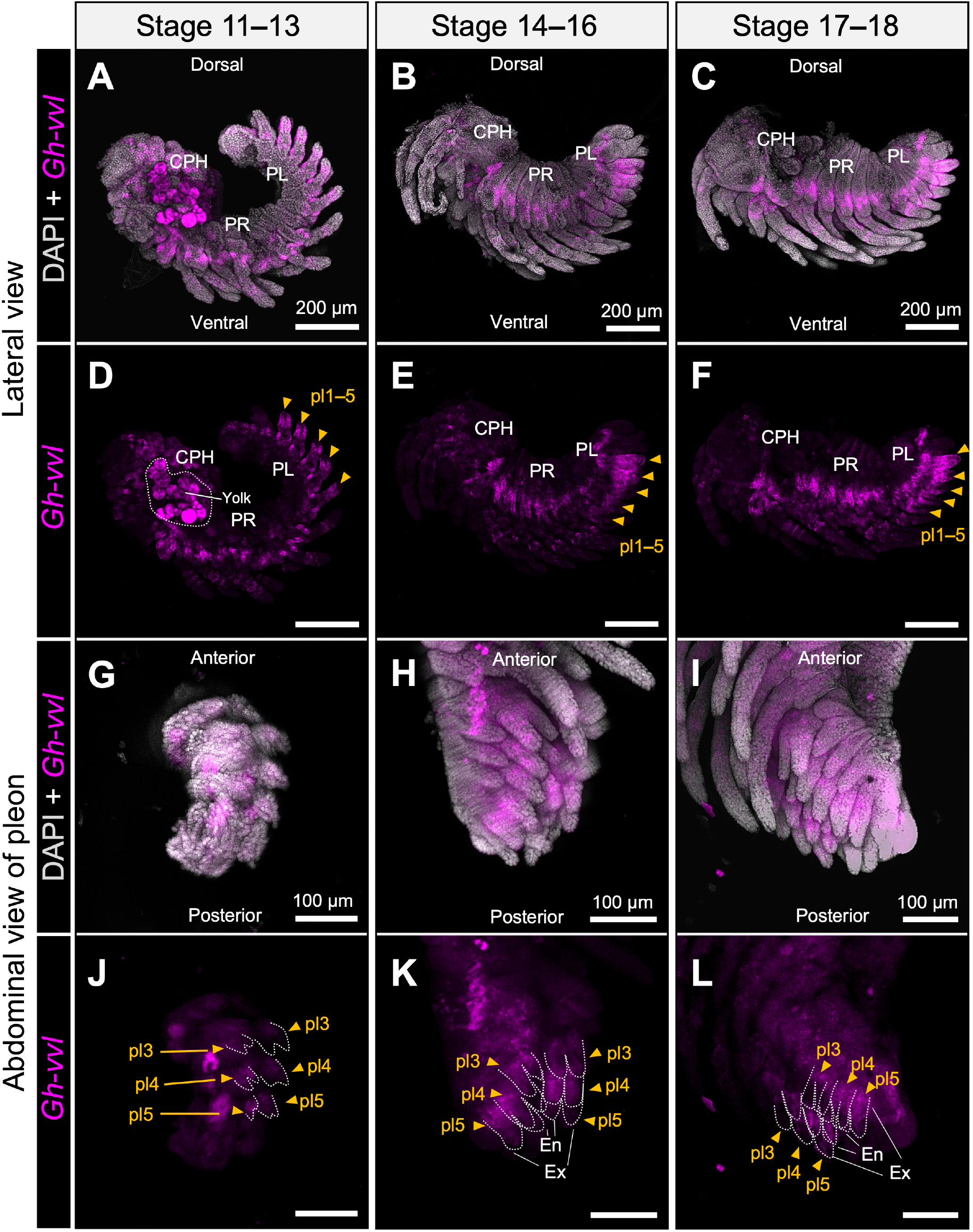
Embryonic expression of *ventral veins lacking* in *Gnorimosphaeroma* cf. *hoestlandti*. HCR localization of *Gh-vvl* across stages 11–13 (A, D, G, J), 14–16 (B, E, H, K) and 17–18 (C, F, I, L). (A–F) Lateral views, with DAPI and *Gh-vvl* merged in A–C and the *Gh-vvl* channel shown in D–F. (G–L) Abdominal views of the pleon, with merged images in G–I and the *Gh-vvl* channel in J–L. *Gh-vvl* signal is magenta and DAPI is gray. Orange arrowheads identify pleopodal expression domains; dotted outlines in J–L indicate selected rami of posterior pleopods. The outlined yolk region in D is not interpreted as gene expression. CPH, cephalon; PR, pereon; PL, pleon; En, endopod; Ex, exopod; pl, pleopod. Scale bars: 200 µm in A–F; 100 µm in G–L.

**Figure 4.**
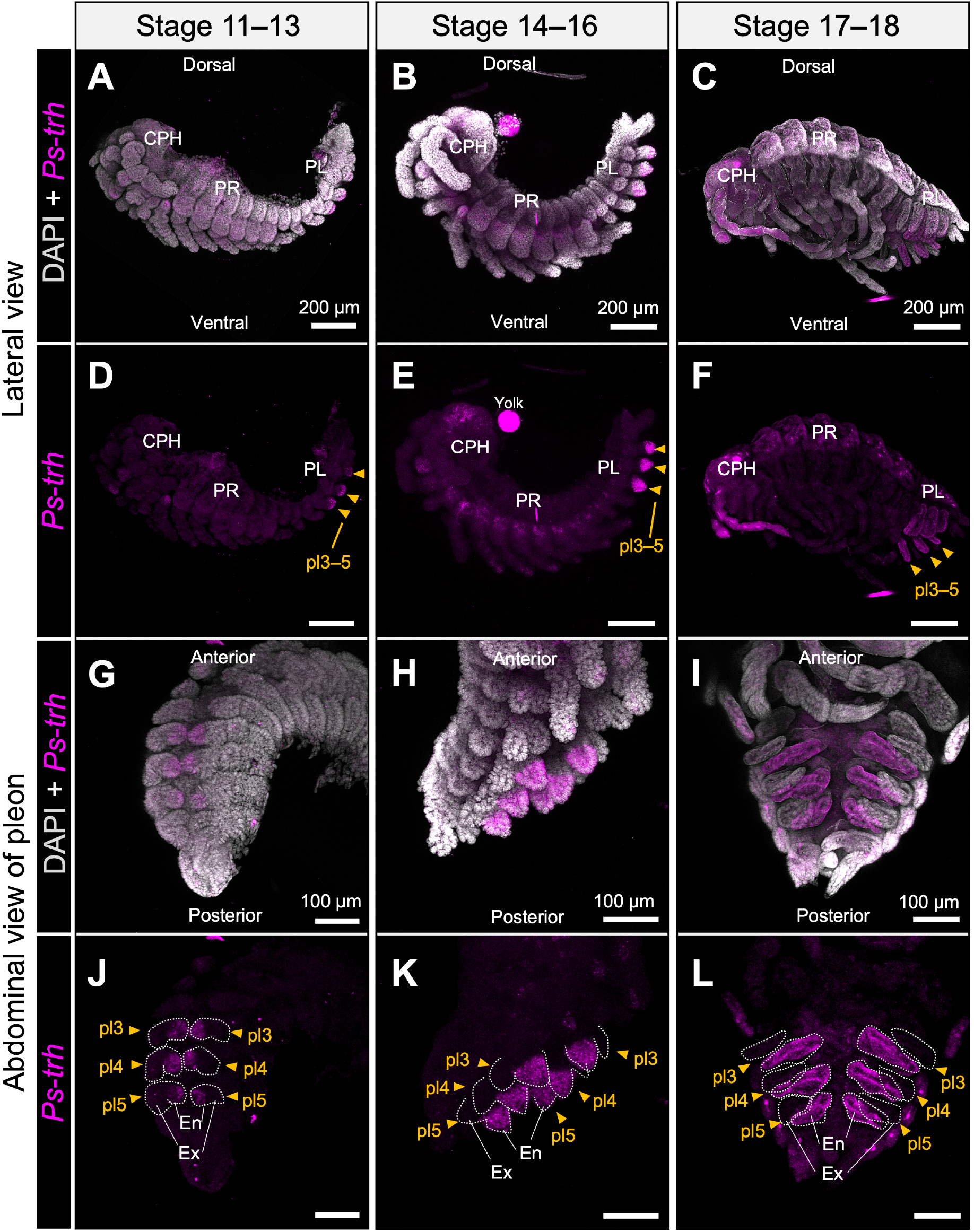
Embryonic expression of *trachealess* in *Porcellio scaber*. HCR localization of *Ps-trh* across stages 11–13 (A, D, G, J), 14–16 (B, E, H, K) and 17–18 (C, F, I, L). (A–F) Lateral views, with DAPI and *Ps-trh* merged in A–C and the *Ps-trh* channel shown in D–F. (G–L) Abdominal views of the pleon, with merged images in G–I and the *Ps-trh* channel in J–L. *Ps-trh* signal is magenta and DAPI is gray. Orange arrowheads identify pleopods 3–5. Dotted outlines indicate selected pleopodal rami, showing the prominent endopodal localization. The structure labeled yolk in E is excluded from expression interpretation. CPH, cephalon; PR, pereon; PL, pleon; En, endopod; Ex, exopod; pl, pleopod. Scale bars: 200 µm in A–F; 100 µm in G–L.

**Figure 5.**
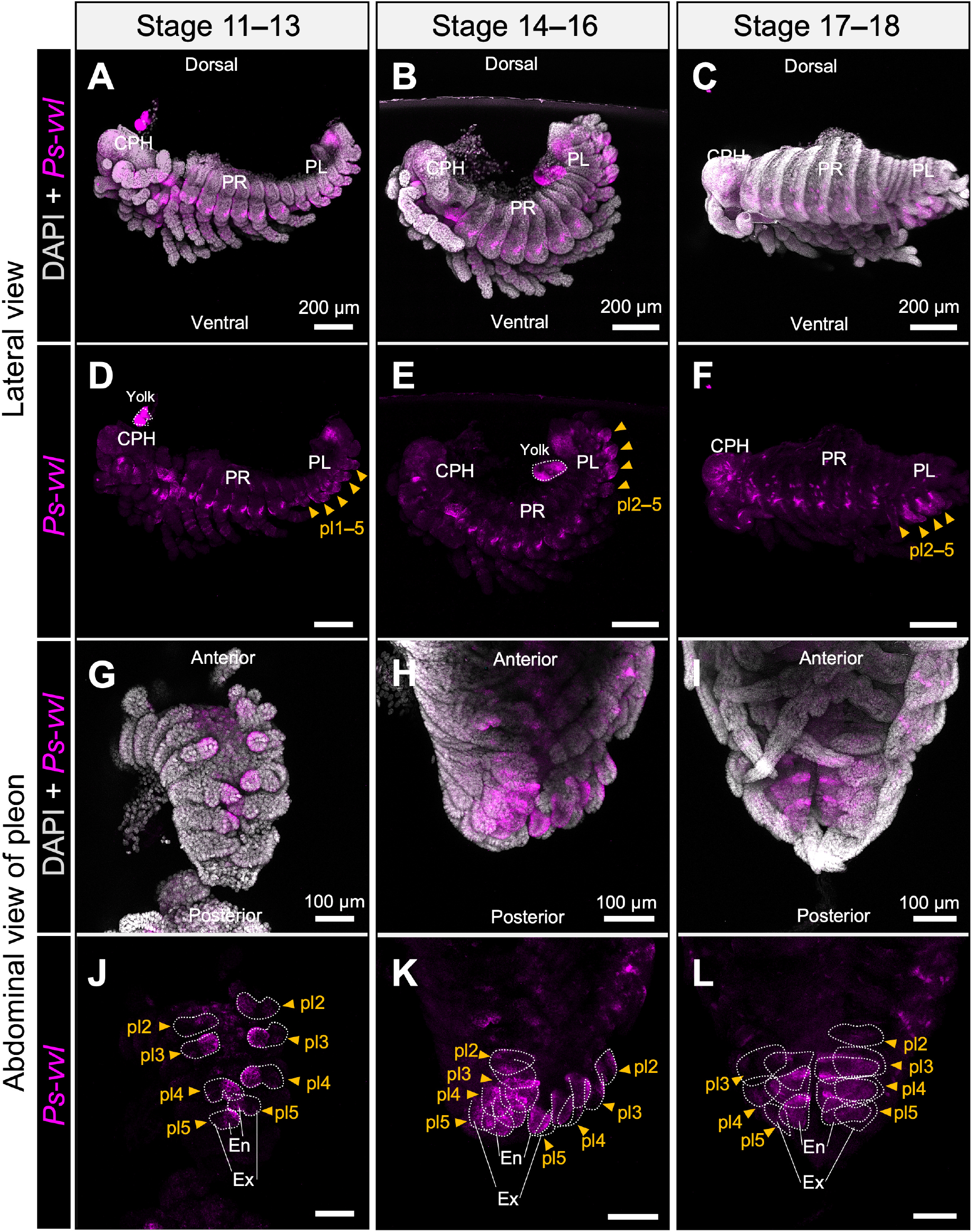
Embryonic expression of ventral veins lacking in Porcellio scaber. HCR localization of *Ps-vvl* across stages 11–13 (A, D, G, J), 14–16 (B, E, H, K) and 17–18 (C, F, I, L). (A–F) Lateral views, with DAPI and *Ps-vvl* merged in A–C and the *Ps-vvl* channel shown in D–F. (G–L) Abdominal views of the pleon, with merged images in G–I and the *Ps-vvl* channel in J–L. *Ps-vvl* signal is magenta and DAPI is gray. Orange arrowheads identify pleopodal domains, including early signal in pleopods 1–5 and later signal in pleopods 2–5. Dotted outlines in J–L indicate selected rami. Regions labeled yolk are excluded from expression interpretation. CPH, cephalon; PR, pereon; PL, pleon; En, endopod; Ex, exopod; pl, pleopod. Scale bars: 200 µm in A–F; 100 µm in G–L.

### *trh* and *vvl* expression includes the gill-bearing rami in the aquatic isopod

We next examined the embryonic expression of *trh* and *vvl* in *G.* cf. *hoestlandti* by HCR. Within the pleon, *Gh-trh* expression became clearly localized to pleopods 4 and 5 during the middle and late developmental stages (Fig. 2, orange arrowheads; Fig. S2). At stages 11–13, pleopodal signals were weak and only faintly detectable (Fig. 2A, D, G, J; Fig. S2G), whereas expression was evident around the bases of cephalic appendages and pereopods (Fig. S2A, D). At stages 14–18, pleopodal expression became more prominent, with a stronger signal in pleopod 5 than in pleopod 4 (Fig. 2E, F; Fig. S2H, I). Abdominal views showed that these expression domains extended into both the endopodal and exopodal regions of the posterior pleopods (Fig. 2H, I, K, L). Additional *Gh-trh* expression was detected in cephalic structures, including the mandible (Fig. S2B, C).

*Gh-vvl* showed a broader expression pattern than *Gh-trh*. At stages 11–13, strong signals were detected in pleopods 1–5, together with repeated expression domains around the bases of appendages and coxal plates (Fig. 3A, D, G, J; Fig. S3A, D, G). Pleopodal expression persisted at later stages, with particularly prominent signals in the posterior pleopods, especially pleopods 4 and 5 (Fig. 3E, F; Fig. S3H, I). Abdominal views showed expression in both endopodal and exopodal regions (Fig. 3K, L). Outside the pleon, *Gh-vvl* was also expressed in cephalic structures, including maxilla 2, and in repeated proximal pereopodal and coxal-plate domains (Fig. S3). Thus, despite their different overall expression patterns, both *Gh-trh* and *Gh-vvl* were expressed in the posterior endopodal and exopodal pleopod regions associated with branchial tissues. *Gh-trh* showed a more restricted posterior pleopodal pattern, whereas *Gh-vvl* exhibited a broader appendage-associated distribution.

### *trh* and *vvl* expression follows the posterior endopodal gills in the terrestrial isopod

We then examined the corresponding expression patterns in *P. scaber*. *Ps-trh* expression was detected in pleopods 3–5 across all three developmental intervals (Fig. 4, arrowheads; Fig. S4). Pleopodal signals were weak at stages 11–13 but became more prominent at subsequent stages (Fig. 4D–F; Fig. S4G–I). As the pleopodal rami differentiated, abdominal views showed that the signals were predominantly localized to the endopods (Fig. 4G–L). By stages 17–18, elongated *Ps-trh* expression domains extended along the posterior endopods (Fig. 4I, L). Outside the pleon, expression was detected around appendage bases and coxal plates, including prominent signals near the maxilliped at early stages and in the antennal region at later stages (Fig. S4A–C).

As in *G.* cf. *hoestlandti, Ps-vvl* showed a broader expression pattern than *Ps-trh* (Fig. 5; Fig. S5). At stages 11–13, *Ps-vvl* expression was observed in pleopods 1–5, whereas at stages 14– 18, it was detected in pleopods 2–5, consistent with the developmental reduction of pleopod 1 (Wolff, 2009) (Fig. 5A–F, arrowheads). Abdominal views showed prominent expression in the posterior endopods, particularly those of pleopods 3–5 (Fig. 5G–L, arrowheads). *Ps-vvl* was also expressed outside the posterior pleopods, including cephalic regions and repeated proximal pereopodal and coxal-plate domains (Fig. S5).

Both *Ps-trh* and *Ps-vvl* were expressed in the posterior endopodal regions associated with gill tissues, although *Ps-vvl* occupied a broader segmental and appendage-associated domain than *Ps-trh*.

### Posterior Hox expression provides a regional context for the gill-associated domains

To place the posterior *trh* and *vvl* expression domains within the regional patterning of the pleon, we examined *abdA* and *AbdB* together with either *trh* or *vvl* by multiplex HCR (Fig. 6; Fig. S6). These comparisons focused on stages 14–16, when the pleopodal expression domains were clearly distinguishable. Negative-control preparations showed no comparable localized signals. Fluorescence in the yolk, hepatopancreatic regions and debris was also observed in controls and was therefore excluded from the interpretation of specific gene expression (Fig. S7).

**Figure 6.**
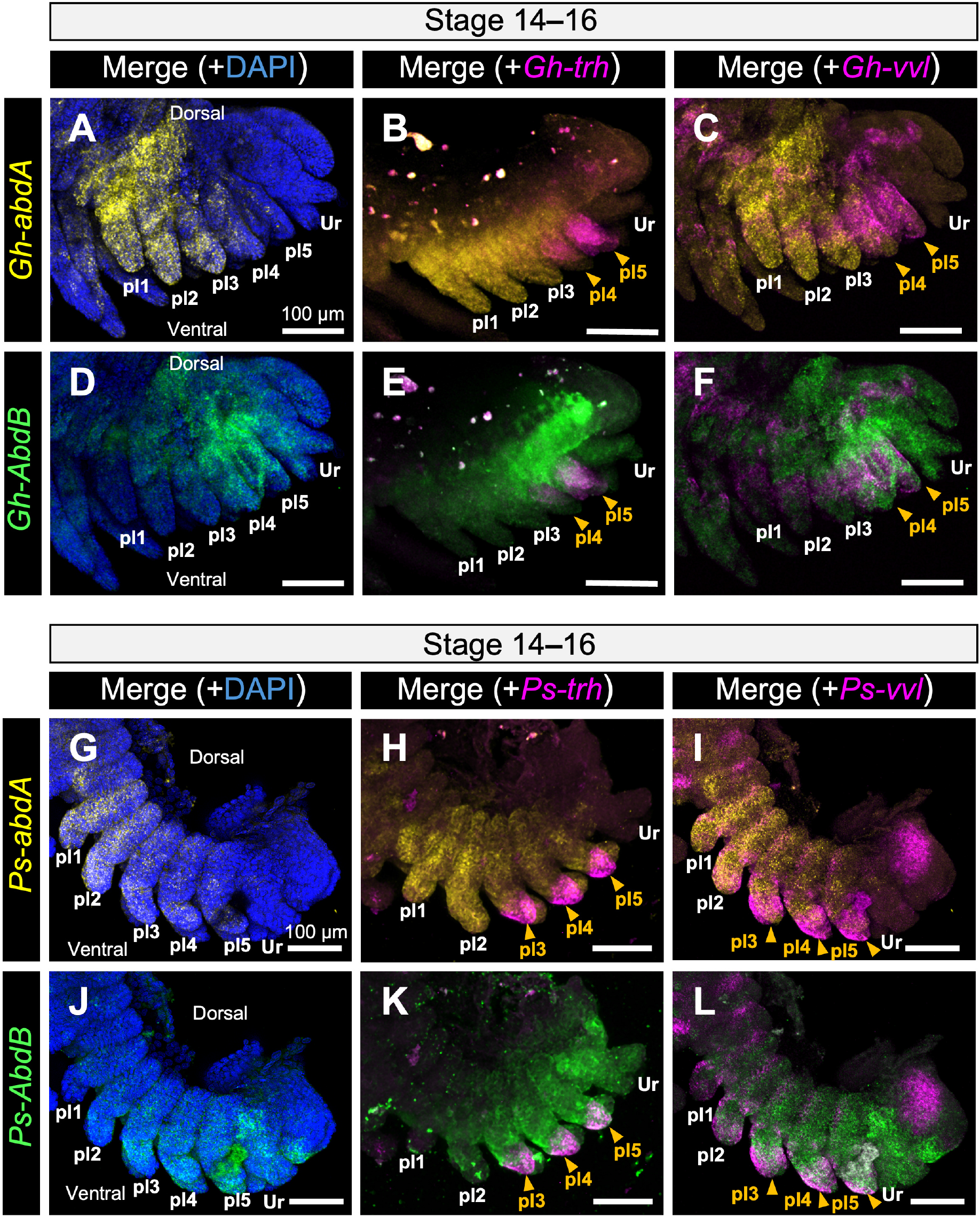
Spatial relationships between posterior Hox expression and pleopodal *trh* or *vvl* expression. Multiplex HCR images of the pleon in *Gnorimosphaeroma* cf. *hoestlandti* (A–F) and *Porcellio scaber* (G–L). (A–C, G–I) *abdA* signal, shown in yellow, merged with DAPI (A, G), *trh* (B, H) or *vvl* (C, I). (D–F, J–L) *AbdB* signal, shown in green, merged with DAPI (D, J), *trh* (E, K) or *vvl* (F, L). DAPI is blue; *trh* and *vvl* signals are magenta. Orange arrowheads identify posterior pleopods with respiratory-associated gene expression. Merged channels locate posterior gill-associated expression within the regional Hox pattern. pl1–pl5, pleopods 1–5; Ur, uropod. Scale bars: 100 µm.

In *G.* cf. *hoestlandti*, *Gh-abdA* expression was most conspicuous in the anterior pleon (Fig. 6A–C; Fig. S6B, F), whereas *Gh-AbdB* expression extended through the posterior pleon and posterior pleopods (Fig. 6D–F; Fig. S6C, G). The localized *Gh-trh* and posterior *Gh-vvl* expression domains in pleopods 4 and 5 spatially overlapped with regions expressing *Gh-AbdB* (Fig. 6A–F, orange arrowheads).

In *P. scaber*, *Ps-abdA* expression extended throughout much of the pleon but was more prominent anteriorly (Fig. 6G–I; Fig. S6J, N). *Ps-AbdB* also showed broad pleonal expression, with particularly prominent signals in the posterior pleopods, where its expression domain spatially overlapped with the *Ps-trh* and *Ps-vvl* domains in pleopods 3–5 (Fig. 6J–L; Fig. S6K, O), consistent with previous observations (Abzhanov & Kaufman, 2000a; Brena et al., 2005).

Although the overall posterior Hox expression patterns were similar between the two species, the anterior extent of the region showing prominent *AbdB* expression differed. In *G.* cf. *hoestlandti*, prominent posterior expression was observed from approximately pleopod 4, whereas in *P. scaber* it extended anteriorly to approximately pleopod 3 (Fig. 6; Fig. S6). This difference approximately paralleled both the anterior extent of localized *trh* expression and the distribution of the posterior gill-bearing pleopods in the two species.

## Discussion

### Pleopodal gill morphology and gill-associated *trh* and *vvl* expression

The histological observations revealed morphological features consistent with branchial tissue in the posterior pleopods of both species (Fig. 1). Crustacean gills are generally characterized by thin cuticles and specialized epithelia associated with hemolymph spaces, which facilitate gas and ion exchange (Freire et al., 2008). In sphaeromatid isopods, pleopods 4 and 5 are generally considered to have branchial functions (Iverson, 1982). Although external branchial folds are not evident in *Gnorimosphaeroma* (Wetzer et al., 2018), our histological observations revealed more extensively folded epithelia and thinner cuticles in pleopods 4 and 5 than in the anterior pleopods, consistent with their proposed branchial function. In *P. scaber*, the posterior pleopodal endopods were covered by thin cuticles, consistent with the morphological characteristics of pleopodal gills previously described in terrestrial isopods (Csonka et al., 2013; Kümmel, 1984; Surbida & Wright, 2001).

Against this anatomical background, the central finding of this study is that *trh* and *vvl* expression occurs in pleopodal regions corresponding to gill-bearing rami in two isopods with different distributions of branchial tissues (Fig. 7A). In the aquatic *G.* cf. *hoestlandti*, prominent posterior expression domains extend into both the endopodal and exopodal regions of pleopods 4 and 5, whereas in the terrestrial *P. scaber*, the prominent posterior expression domains are predominantly associated with the endopods of pleopods 3–5. Thus, the spatial correspondence between *trh* and *vvl* expression and gill-bearing pleopodal regions is observed in both species despite differences in the distribution of branchial tissues.

**Figure 7.**
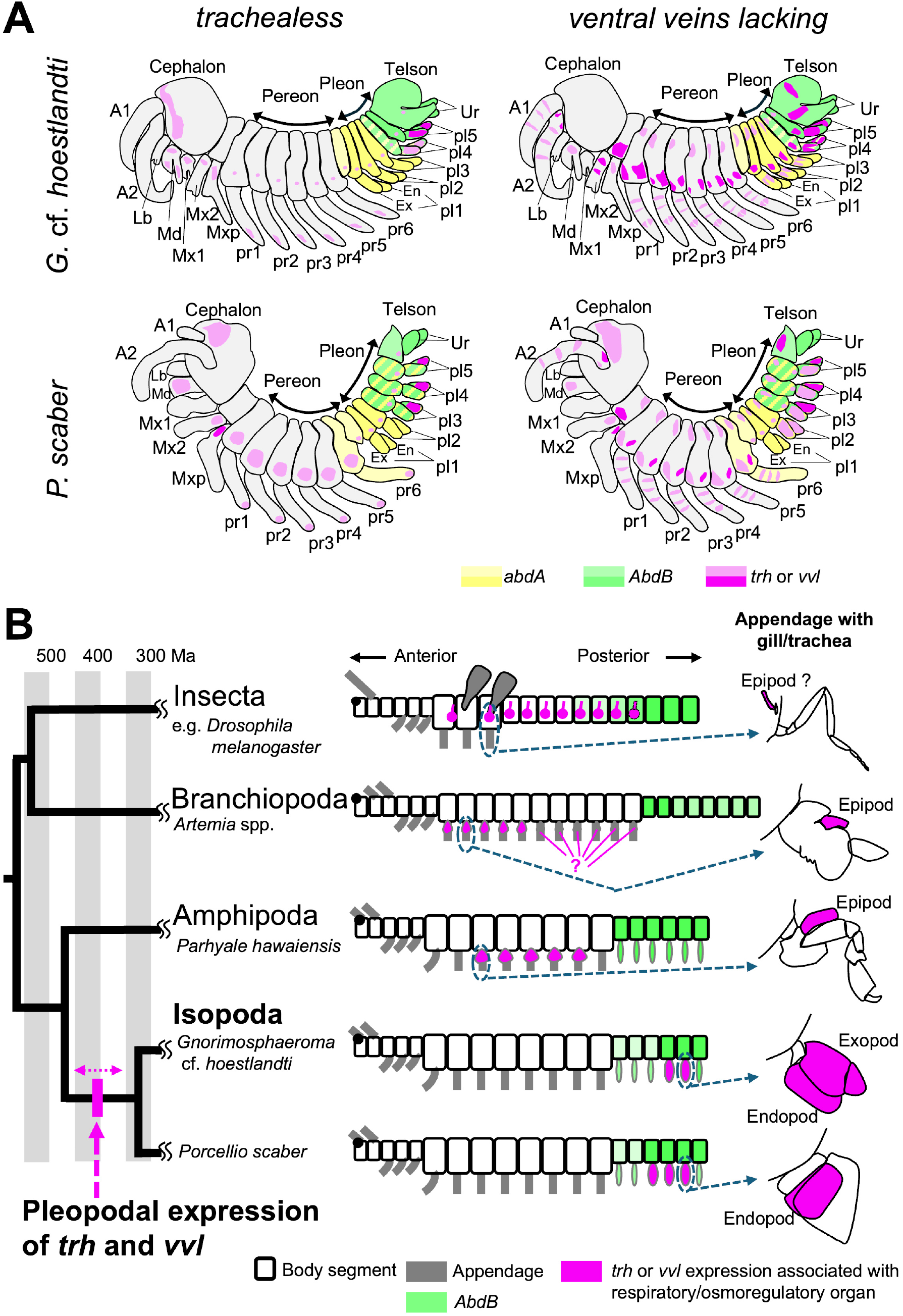
Heterotopic expression of respiratory-associated genes. (A) Schematic summary of *trh* and *vvl* expression in the two focal isopod embryos, shown against the regional distributions of *abdA* and *AbdB*. Yellow indicates *abdA*, green *AbdB*, and magenta *trh* or *vvl*. The figure shows the expression patterns observed in mid-stage embryos (stages 14–16). Darker and lighter shades indicate qualitatively stronger and weaker HCR signals, respectively. They do not represent quantitative differences in expression levels across genes or species. (B) Comparison of pancrustacean body plans and respiratory-associated expression domains within a simplified, time-calibrated phylogenetic framework. Divergence times among pancrustacean lineages are based on Bernot et al. (2023), whereas those within Isopoda are based on Thomas Thorpe (2024). Gray and white bands each represent 50-million-year intervals, with the labeled boundaries corresponding to 300, 400, and 500 million years ago (Ma). Published *trh* and/or *vvl* expression in *Artemia* and *Parhyale* is associated with thoracic epipods, whereas the focal isopod domains occur in pleopodal rami. The dashed magenta arrow represents hypothesized heterotopic deployment during pleopodal gill evolution, with its position approximately corresponding to the divergence of the isopod lineage. The insect comparison concerns tracheal tissue and its proposed evolutionary relationship to epipodal structures. Comparative gene-expression information is based on previous studies (Franch-Marro et al., 2006; Hanna & Popadić, 2020; Mitchell & Crews, 2002; Wang et al., 2012). A1 and A2, first and second antennae; Lb, labrum; Md, mandible; Mx1 and Mx2, first and second maxillae; Mxp, maxilliped; pr, pereopod; pl, pleopod; En, endopod; Ex, exopod; Ur, uropod. Body-plan and appendage schematics are not to scale; the phylogeny is simplified and divergence times are approximate.

The two genes nevertheless show distinct spatial characteristics within the pleon. *trh* forms relatively restricted posterior domains, concentrated in pleopods 4 and 5 in *G.* cf. *hoestlandti* and pleopods 3–5 in *P. scaber*, whereas *vvl* is expressed more broadly across pleopods and other appendage-associated regions. Previous studies reported *vvl* expression in the posterior pleopods of *P. scaber* (Abzhanov & Kaufman, 2000b; Wolff, 2009), and adult transcriptomic analyses showed higher expression of both *trh* and *vvl* in endopods than in exopods (Inui et al., 2025). Our developmental series extends these observations by showing that *trh* and *vvl* signals are already detectable in posterior pleopods at stages 11–13 and that their spatial association with gill-bearing regions becomes more apparent as the pleopodal rami differentiate. The broader expression of *vvl*, however, suggests that its developmental functions extend beyond the pleopodal gills. In *Drosophila*, Vvl functions not only in tracheal development but also in the differentiation of other cell types, including midline glia (Anderson et al., 1995), consistent with a broader developmental regulatory role. Likewise, the repeated *vvl* expression domains observed in cephalic and thoracic appendage-associated regions in the present study may reflect additional developmental roles in these regions, although their functions remain unknown. Thus, although the posterior *vvl* domains consistently include gill-bearing regions, this association should be interpreted within a broader, potentially pleiotropic developmental role. The same interpretation applies to *trh*, which also shows expression outside the pleopods.

Notably, the expression of both *trh* and *vvl* in pleopodal gill regions does not necessarily extend to all respiratory structures. In our previous analysis of *P. scaber*, neither *trh* nor *vvl* showed preferential expression in the derived exopodal lungs, despite their higher expression in the pleopodal endopods (Inui et al., 2025). Together with the present embryonic observations, these findings suggest that the expression patterns of *trh* and *vvl* are more closely associated with the retained pleopodal gill system than with respiratory structures in general.

### Posterior Hox patterning provides a candidate regional context for gill differentiation

The posterior Hox expression patterns provide a potential axial context for the gill-associated *trh* and *vvl* domains. Hox genes have well-established roles in specifying regional and appendage identities in crustaceans: shifts in Hox expression boundaries correlate with appendage diversification (Averof & Akam, 1995), and functional analyses in *Parhyale* have demonstrated roles for Hox genes in appendage specification, including a requirement for Ubx in thoracic gill development (Averof & Akam, 1995; Martin et al., 2016). In *P. scaber*, *AbdB* was previously reported to become localized to the endopods of pleopods 3–5 during late embryogenesis, raising the possibility that this gene contributes to their respiratory specialization (Brena et al., 2005).

Our comparison extends this spatial relationship to an aquatic isopod. In both species, *abdA* expression was more prominent in the anterior pleon, whereas *AbdB* expression extended into the posterior pleopods. The posterior *trh* and *vvl* expression domains spatially overlapped with these *AbdB*-prominent regions. Notably, the anterior extent of prominent *AbdB* expression approximately corresponded to that of the localized *trh* domains: both extended anteriorly to approximately pleopod 4 in *G.* cf. *hoestlandti* and pleopod 3 in *P. scaber*. This spatial correspondence suggests a possible association between posterior Hox regionalization and the species-specific distribution of gill-bearing pleopodal regions. These observations raise the possibility that *AbdB* contributes to defining the pleopodal regions in which gill differentiation occurs. Although the spatial correspondence alone does not establish a regulatory relationship, functional analyses of *AbdB* and its effects on *trh* and *vvl* expression and pleopodal morphology would help test this hypothesis.

### Heterotopic deployment of respiratory-associated genes during pleopodal gill evolution

Comparisons with other crustaceans provide an evolutionary context for the pleopodal expression patterns observed in this study (Fig. 7B). Thoracic epipods or epipod-associated structures form gills in many crustacean lineages (Boxshall & Jaume, 2009). In *Artemia* and *Parhyale*, *trh* and *vvl* homologues are expressed in developing thoracic epipods, including epipodal gills (Franch-Marro et al., 2006; Mitchell & Crews, 2002). By contrast, in the two isopods examined here, *trh* and *vvl* show prominent expression in gill-bearing pleopodal regions (Fig. 7A, B). This contrast involves two anatomical levels: the body axis (thoracic versus abdominal appendages) and the appendage structures bearing gill-associated expression (epipods versus endopods and exopods). At a broader pancrustacean scale, *trh* and *vvl* also contribute to insect tracheal development, although the evolutionary relationship between the tracheal system and ancestral crustacean epipodal structures remains unresolved (Franch-Marro et al., 2006; Hanna & Popadić, 2020).

These observations are consistent with the hypothesis that pleopodal gill evolution involved heterotopic deployment of developmental regulatory genes already associated with respiratory or osmoregulatory tissues. This hypothesis concerns the spatial redeployment of developmental regulators rather than structural homology between pleopodal rami and thoracic epipods. Shared expression of developmental regulators can occur in non-homologous structures, and previous studies of *P. scaber* have cautioned against inferring structural homology from *vvl* or *AbdB* expression alone (Brena et al., 2005; Wolff, 2009). Furthermore, both *trh* and particularly *vvl* showed repeated expression around proximal thoracic appendage and coxal-plate regions in the focal isopods (Figs. S2–S5). Given the proximal position of epipods in other crustaceans, these thoracic expression domains raise the possibility that pleopodal expression was added while elements of an ancestral proximal appendage expression program were retained, rather than simply replacing a thoracic expression domain. However, the homology of these thoracic expression domains remains uncertain.

The time-calibrated phylogeny provides a temporal framework for interpreting the evolution of these expression patterns (Fig. 7B). Respiratory-associated expression of *trh* and *vvl* in distantly related lineages, including *Artemia* and hexapods, whose divergence dates back more than 550 million years ago (Ma), suggests that their involvement in respiratory development may have ancient evolutionary origins (Bernot et al., 2023). In contrast, expression in gill-bearing pleopodal rami has so far been reported only in isopods, whereas respiratory-associated expression in other examined crustaceans occurs in thoracic epipods. This phylogenetic distribution supports the hypothesis that pleopodal expression represents a derived feature acquired in the isopod lineage after its divergence from other peracarids. Given the widespread occurrence of pleopodal gills in Isopoda, this heterotopic deployment may have preceded the earliest divergences within the group, estimated at approximately 424 Ma (Thomas Thorpe, 2024). The different branchial arrangements and associated expression patterns of *G.* cf. *hoestlandti* (Sphaeromatidae) and *P. scaber* (Porcellionidae, Oniscidea), whose lineages diverged approximately 338 Ma (Thomas Thorpe, 2024), further suggest that the spatial distribution of these genes changed in at least one lineage after their divergence.

A broader phylogenetic comparison will be particularly informative for clarifying the evolutionary origin and subsequent diversification of pleopodal expression. Respiratory epipods occupy different serial positions among malacostracan lineages, ranging from multiple thoracopods to more restricted maxillipedal distributions (Boxshall & Jaume, 2009; Tan & Monteiro, 2025). Comparing *trh* and *vvl* expression in these taxa, including closely related peracarids such as cumaceans, and in additional aquatic isopods with different endopodal and exopodal gill arrangements would help establish when pleopodal expression arose and whether subsequent changes in its segmental or ramal distribution accompanied the diversification of isopod gills.

## Conclusion

Our results show that embryonic *trh* and *vvl* expression corresponds to pleopodal gill-bearing regions in aquatic and terrestrial isopods with different branchial architectures. In both species, these posterior pleopodal expression domains spatially overlap with *AbdB*-expressing regions, while differences in their segmental and ramal distributions parallel differences in gill organization. Comparison with the thoracic epipodal expression of *trh* and *vvl* reported in other crustaceans suggests that pleopodal gill evolution may have involved the heterotopic deployment of developmental regulators with ancient associations to respiratory tissues. The phylogenetic framework further suggests that this deployment may have originated early in isopod evolution, preceding the transition to various habitats, including terrestrial environments. Functional analyses of *trh* and *vvl* in crustaceans and broader phylogenetic comparisons will be required to determine how these spatial expression patterns are generated and whether their evolutionary modification contributed causally to the diversification of crustacean gill positions.

## Materials and Methods

### Animals

Terrestrial isopods, *Porcellio scaber*, were collected from soil on the campuses of Misaki Marine Biological Station, The University of Tokyo, Koajiro, Misaki, Kanagawa, Japan and the National Institute of Genetics, Mishima, Shizuoka, Japan. Some individuals were maintained under laboratory conditions as described previously (Inui et al., 2022). Aquatic isopods, *Gnorimosphaeroma* cf. *hoestlandti*, were collected from Arai-hama Beach, Misaki, Kanagawa Prefecture, and Wakano-ura Beach, Shimoda, Shizuoka Prefecture, Japan. *Gnorimosphaeroma* specimens were identified based on previous descriptions (Kim & Kwon, 1985; Nunomura & Shimomura, 2016). Although the collected specimens closely matched the characteristics of *G. hoestlandti*, slight differences were observed in aspects such as the arrangement of setae on the first pereopod. Therefore, this species was treated as *G.* cf. *hoestlandti* in the present study. Some individuals were maintained in a 30-cm aquarium containing aerated seawater and fed brown algae, such as *Sargassum* spp.

For both species, embryos were obtained from the marsupia of ovigerous females. Embryonic stages were assigned according to the morphological staging system established for *P. scaber* (Wolff, 2009). Because the major morphological features of embryonic development in *G.* cf. *hoestlandti* generally corresponded to those described for *P. scaber*, the same staging criteria were applied to both species (Fig. S1).

Embryos at stages 11–18, when the appendages were clearly formed, were used for expression analyses. They were grouped into three developmental intervals: early (stages 11–13), middle (stages 14–16), and late (stages 17–18). The chorion, vitelline membrane, yolk, and hepatopancreas were removed as thoroughly as possible using a tungsten needle. Embryos were then fixed in 4% paraformaldehyde in 1× phosphate-buffered saline (PBS) for approximately 30 min. After fixation, embryos were washed in PBS, dehydrated in methanol, and stored at −20°C until use.

### Morphological observations

Pleopodal tissues of the focal species were examined histologically to identify respiratory structures. Adult pleopods of both species were observed using a stereomicroscope (SZX16; Olympus, Tokyo, Japan) equipped with a digital camera (DP50; Olympus). Sagittal sections of the abdomen were prepared and imaged as described previously (Inui et al., 2022).

### Identification of orthologous sequences

Candidate orthologues of *trh*, *vvl*, *abdA*, and *AbdB* were identified from transcriptome assemblies of the two focal species using sequence-similarity searches and phylogenetic analyses. For *P. scaber*, transcriptome data were obtained from the previously published assembly (BioProject accession no. PRJDB20244; Inui et al., 2025).

For *G.* cf. *hoestlandti*, a transcriptome assembly was newly generated in this study. Total RNA was extracted from pooled samples representing various developmental stages using ISOGEN (Nippon Gene, Tokyo, Japan). The extracted RNA was treated with DNase I (Thermo Fisher Scientific, MA, USA) and purified using RNA Clean XP beads (Beckman Coulter Genomics, MA, USA). Library preparation and sequencing were performed by Genome-Lead Inc. (Kagawa, Japan). Libraries were prepared using the MGIEasy RNA Directional Library Prep Set (MGI Tech, Guangdong, China) with the KAPA mRNA Capture Kit (Kapa Biosystems, Boston, MA, USA). DNA nanoballs were generated using a DNBSEQ-T7RS High-Throughput Sequencing Set (MGI Tech) and sequenced on a DNBSEQ-T7RS platform using a 2 × 150-bp paired-end protocol. *De novo* assembly was performed as described previously (Inui et al., 2025). The raw sequencing reads and transcriptome assembly were deposited in the DDBJ databases under accession nos. DRR1079627 (Sequence Read Archive) and ICXZ01000001– ICXZ01458511 (Transcriptome Shotgun Assembly), respectively. The associated BioProject accession number is PRJDB42836.

To identify orthologues of *trh*, *vvl*, *abdA*, and *AbdB*, the corresponding *Drosophila melanogaster* protein sequences were used as queries in tBLASTn searches against the transcriptome assemblies of both species. The top candidate hits were translated and aligned with selected pancrustacean protein sequences using MAFFT v7.526 (Katoh & Standley, 2013) (Tables S1 and S2). For some genes, additional highly similar transcript sequences were present in the assemblies and were regarded as putative transcript isoforms of the identified orthologues. The top-hit sequences were used as representative sequences for the subsequent phylogenetic analyses. Poorly aligned regions were trimmed using trimAl v1.5 with the -gappyout option (Capella-Gutiérrez et al., 2009). Maximum-likelihood phylogenetic trees were inferred using IQ-TREE v3.0.1 (Wong et al., 2026), with the best-fit substitution models selected using ModelFinder (Kalyaanamoorthy et al., 2017). Branch support was assessed using 1,000 ultrafast bootstrap replicates. Phylogenetic trees were edited and visualized using iTOL (Letunic & Bork, 2007). Their placements in gene trees supported the gene assignments (Fig. S8).

### *In situ* hybridization chain reaction

Transcript localization was examined by whole-mount *in situ* hybridization chain reaction (HCR) using the split-initiator probe framework (Choi et al., 2018; Tsuneoka & Funato, 2020). DNA probe pairs were designed using HCR 3.0 Probe Maker (Kuehn et al., 2022) to target coding and untranslated regions of the focal transcripts. For genes with multiple highly similar isoform-like transcripts, the probe sets were not designed to distinguish among these sequences and were expected to detect them collectively. After probe design, BLAST searches against the corresponding transcriptome assemblies were used to assess potential off-target hybridization. For each probe pair, highly similar matches were restricted to the target candidate gene and the closely related sequences regarded as its transcript isoforms. Probe oligonucleotides were synthesized by Integrated DNA Technologies (IA, USA). Probe sequences for the focal genes are listed in Tables S3–S10.

HCR was performed using ISHpalette™ (Nepa Gene, Chiba, Japan), following the manufacturer’s instructions (Tsuneoka & Funato, 2020) with the modifications described below. To improve probe penetration, late-stage embryos with developing cuticles (stages 17–18) were sonicated for several seconds before hybridization, as described previously (Inui et al., 2025). Multiplex HCR was used to detect *abdA* and *AbdB* together with either *trh* or *vvl*. Nuclei were counterstained with DAPI to provide an anatomical reference for localizing expression within developing appendages. The samples were mounted on a glass-bottom dish, and observed using a confocal microscope (FV3000; Olympus, Tokyo, Japan) with laser excitations of 405 nm (DAPI), 488 nm (*abdA* with SaraFluor 488), 561 nm (*trh* or *vvl* with ATTO 550), and 640 nm (*AbdB* with Cyanine5). Images were processed with software FV31S-SW (Olympus, Tokyo, Japan) and ImageJ (Schneider et al., 2012). At least 5 embryos were examined for each gene/stage combination. Negative-control HCR experiments were performed using only one oligonucleotide of each split-initiator probe pair.

## Abbreviations

abdA: abdominal-A AbdB Abdominal-B
BLAST: Basic Local Alignment Search Tool
CDS: coding sequence
DAPI: 4′,6-diamidino-2-phenylindole
DDBJ: DNA Data Bank of Japan
HCR: hybridization chain reaction
iTOL: Interactive Tree Of Life
MAFFT: Multiple Alignment using Fast Fourier Transform
PBS: phosphate-buffered saline
trh: trachealess
UTR: untranslated region
vvl: ventral veins lacking

## Declarations

### Ethics approval and consent to participate

Ethics approval was not required for this study involving isopod crustaceans.

### Consent for publication

Not applicable.

### Availability of data and materials

Raw sequencing reads and the transcriptome assembly of *G.* cf. *hoestlandti* have been deposited in DDBJ under BioProject no. PRJDB42836. The previously published *P. scaber* transcriptome is available under BioProject no. PRJDB20244. Accession numbers of the candidate genes are listed in Table S1.

### Competing Interests

The authors declare no competing interests.

### Funding

This study was supported by JSPS KAKENHI Grant Numbers 25KJ0386 and 26K18362 from the Ministry of Education, Culture, Sports, Science and Technology and "Strategic Research Projects" grant from ROIS (Research Organization of Information and Systems).

### Authors’ Contributions

NI and TM designed the study. NI collected embryos from the study species. NI and AY performed transcriptomic and orthologue analyses. NI and KF performed HCR analysis. NI wrote the manuscript with input from all co-authors. All authors approved the final version of the manuscript.

## Acknowledgments

We would like to thank the staff and students of Misaki Marine Biological Station for their assistance in collecting *Gnorimosphaeroma* cf. *hoestlandti*. We also thank Shin-ya Miyagishima for providing access to laboratory facilities. Computations were partly performed using the NIG supercomputer system (National Institute of Genetics at Research Organization of Information and Systems, Japan). ChatGPT (OpenAI) was used to assist with English-language proofreading and improving the clarity of the manuscript. All suggested changes were reviewed by the authors, who take full responsibility for the final content.

## Notes

### Competing Interest Statement

The authors have declared no competing interest.

