## Supplementary figures for "Heterotopic expression of respiratory-associated genes suggests a developmental basis for pleopodal gill evolution in isopod crustaceans"

**for**

### List of Supplementary Materials:

Supplementary Tables S1–S10 (Excel files)
Supplementary Figs. S1–S8 (this document)

### Supplementary Figures

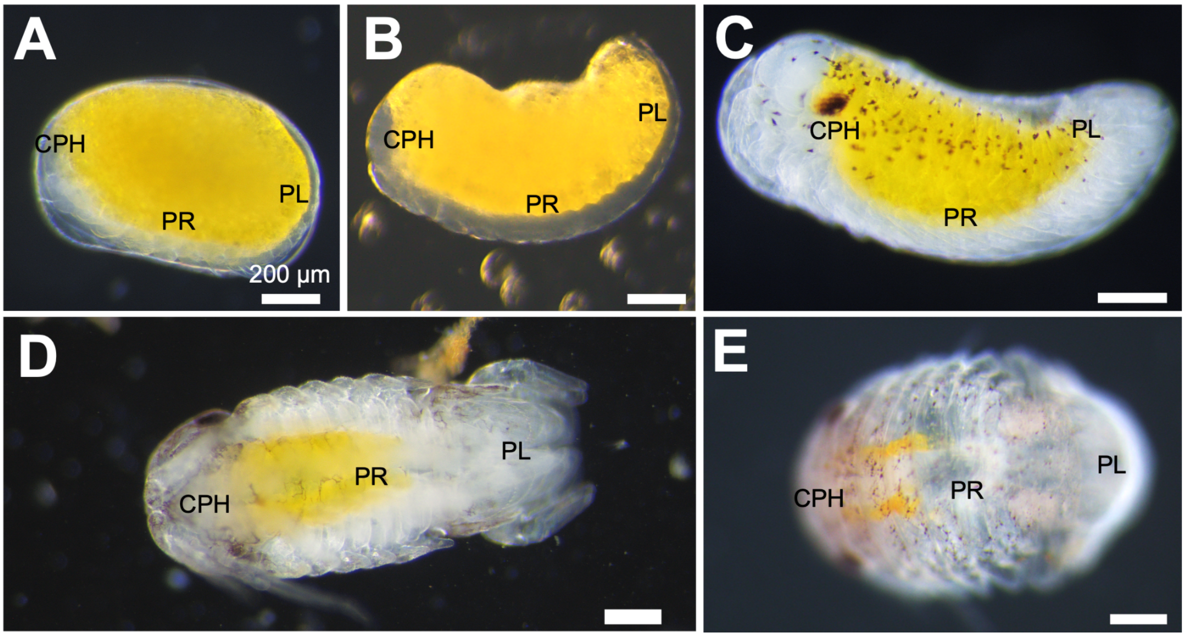


**Figure S1. Developmental stages of *Gnorimosphaeroma* cf. *hoestlandti***

Developmental appearances of living embryos of *G.* cf. *hoestlandti.* (A) Early stage (Stage 11–13). (B) Middle stage (Stage 14–16). (C) Late stage (Stage 17–18). (D) Hatching embryo (first manca stage). (E) Manca (second manca stage). CPH, cephalon; PR, pereon; PL, pleon. Scale bars: 200 µm.

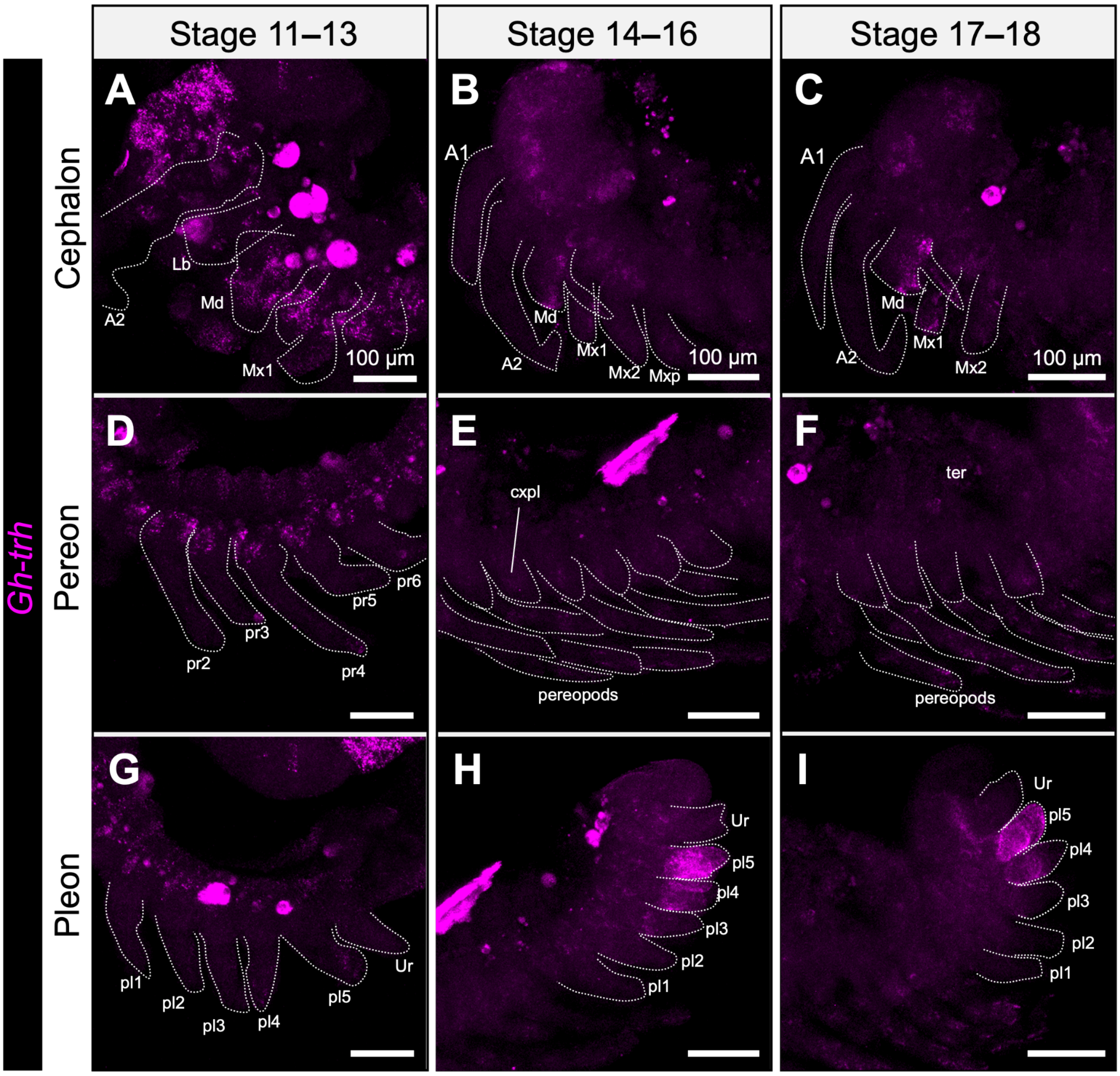


**Figure S2. Regional views of *Gh-trh* expression**

Enlarged lateral views of HCR *Gh-trh* labeling in *Gnorimosphaeroma* cf. *hoestlandti*. Columns show stages 11–13 (A, D, G), 14–16 (B, E, H) and 17–18 (C, F, I); rows show the cephalon (A–C), pereon (D–F) and pleon (G–I). Magenta indicates the *Gh-trh* channel. Dotted lines delimit selected appendages. Yolk-associated fluorescence and the debris identified in Fig. 2 should not be interpreted as additional expression domains. A1 and A2, first and second antennae; Lb, labrum; Md, mandible; Mx1 and Mx2, first and second maxillae; Mxp, maxilliped; cxpl, coxal plate; ter, tergite; pr, pereopod; pl, pleopod; Ur, uropod. Scale bars: 100 µm.

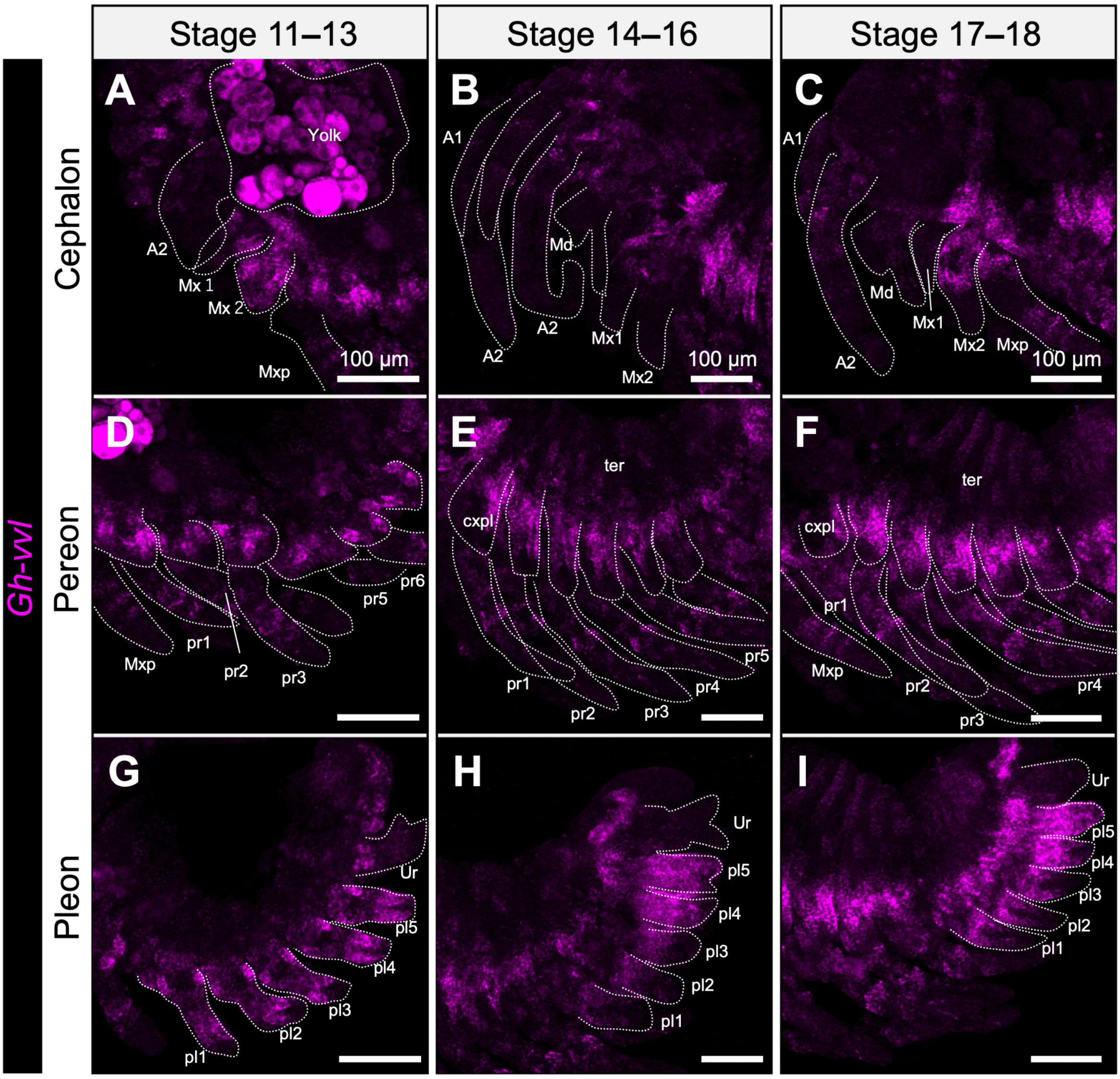


**Figure S3. Regional views of *Gh-vvl* expression**

Enlarged lateral views of HCR *Gh-vvl* labeling in *Gnorimosphaeroma* cf. *hoestlandti*. Columns show stages 11–13 (A, D, G), 14–16 (B, E, H) and 17–18 (C, F, I); rows show the cephalon (A–C), pereon (D–F) and pleon (G–I). Magenta indicates the *Gh-vvl* channel. Dotted lines delimit selected appendages, and yolk-associated fluorescence is identified in A. A1 and A2, first and second antennae; Md, mandible; Mx1 and Mx2, first and second maxillae; Mxp, maxilliped; cxpl, coxal plate; ter, tergite; pr, pereopod; pl, pleopod; Ur, uropod. Scale bars: 100 µm.

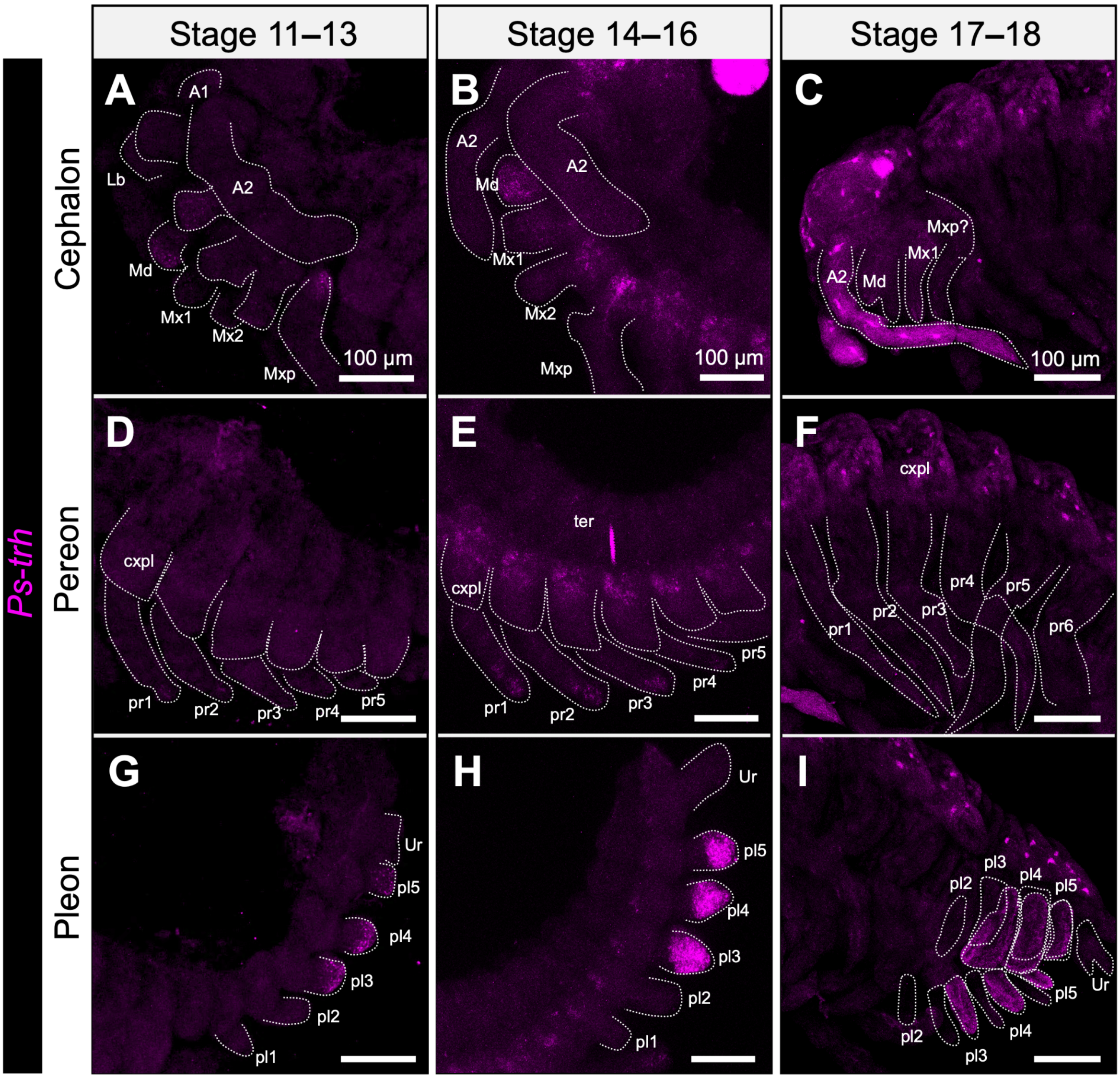


**Figure S4. Regional views of *Ps-trh* expression**

Enlarged views of HCR *Ps-trh* labeling in *Porcellio scaber*. Columns show stages 11–13 (A, D, G), 14–16 (B, E, H) and 17–18 (C, F, I); rows show the cephalon (A–C), pereon (D–F) and pleon (G–I). Magenta indicates the *Ps-trh* channel. Dotted lines delimit selected appendages. A1 and A2, first and second antennae; Lb, labrum; Md, mandible; Mx1 and Mx2, first and second maxillae; Mxp, maxilliped; cxpl, coxal plate; ter, tergite; pr, pereopod; pl, pleopod; Ur, uropod. Scale bars: 100 μm.

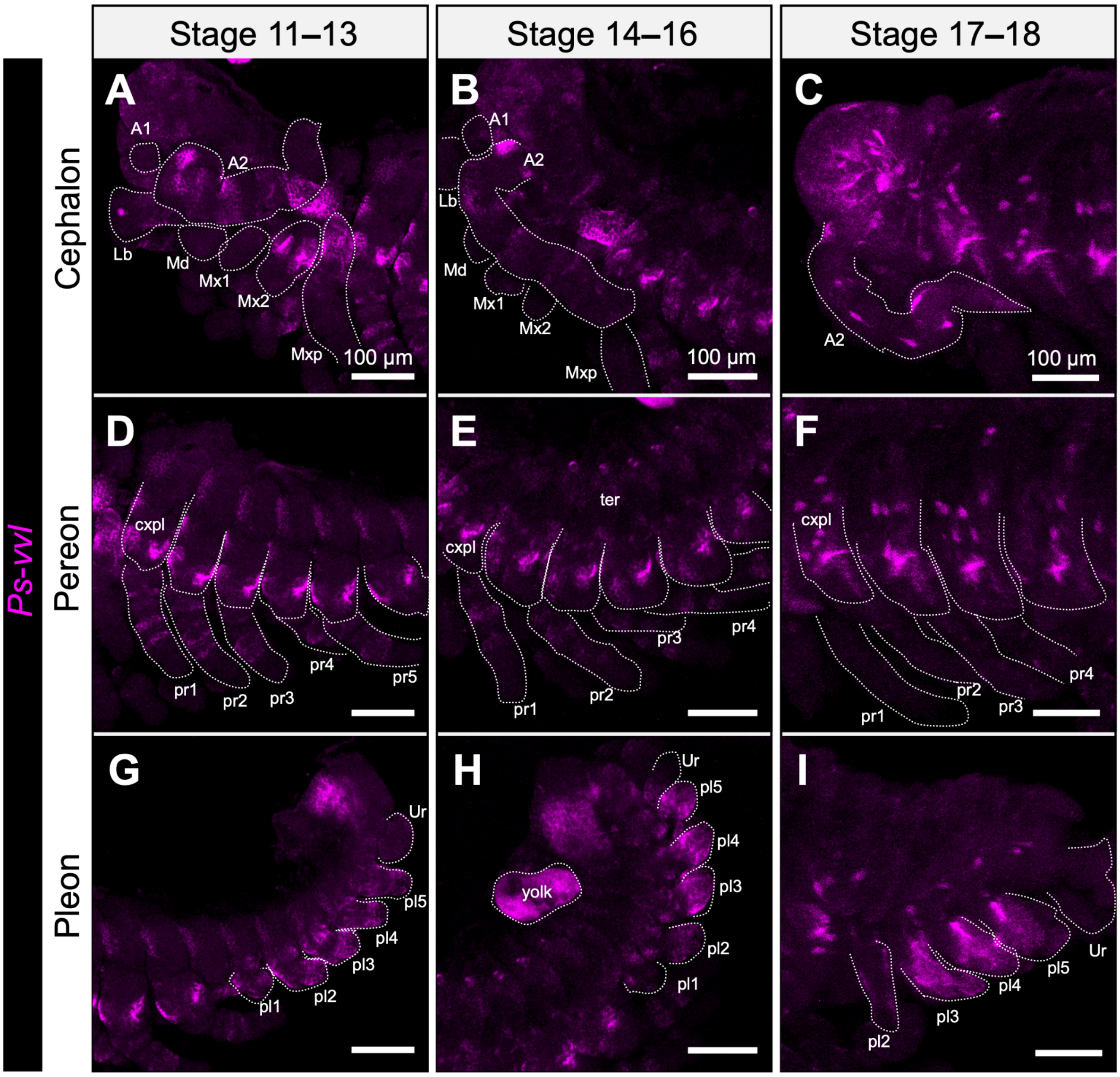


**Figure S5. Regional views of *Ps-vvl* expression**

Enlarged views of HCR *Ps-vvl* labeling in *Porcellio scaber*. Columns show stages 11–13 (A, D, G), 14–16 (B, E, H) and 17–18 (C, F, I); rows show the cephalon (A–C), pereon (D–F) and pleon (G–I). Magenta indicates the *Ps-vvl* channel. Dotted lines delimit selected appendages; the yolk region is labeled in H. A1 and A2, first and second antennae; Lb, labrum; Md, mandible; Mx1 and Mx2, first and second maxillae; Mxp, maxilliped; cxpl, coxal plate; ter, tergite; pr, pereopod; pl, pleopod; Ur, uropod. Scale bars: 100 µm.

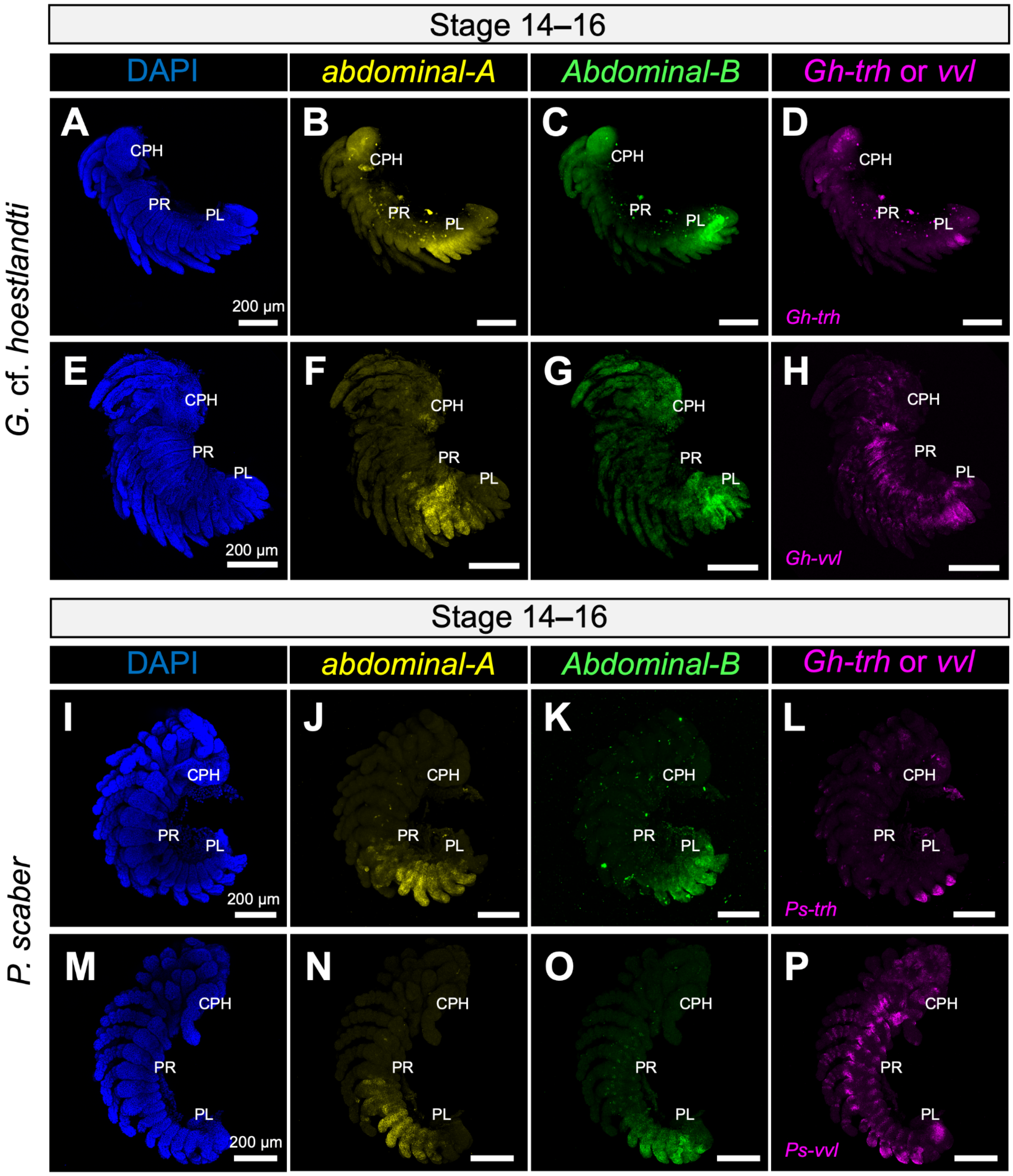


**Figure S6. Whole-embryo views of posterior Hox and respiratory-associated gene expression**

Separate channels from multiplex HCR preparations of *Gnorimosphaeroma* cf. *hoestlandti* (A–H) and *Porcellio scaber* (I–P). Columns show DAPI (A, E, I, M; blue), *abdA* (B, F, J, N; yellow), *AbdB* (C, G, K, O; green), and either *trh* (D, L) or *vvl* (H, P), shown in magenta. These views provide the body-wide context for the pleopodal comparisons. Scale bars: 200 µm.

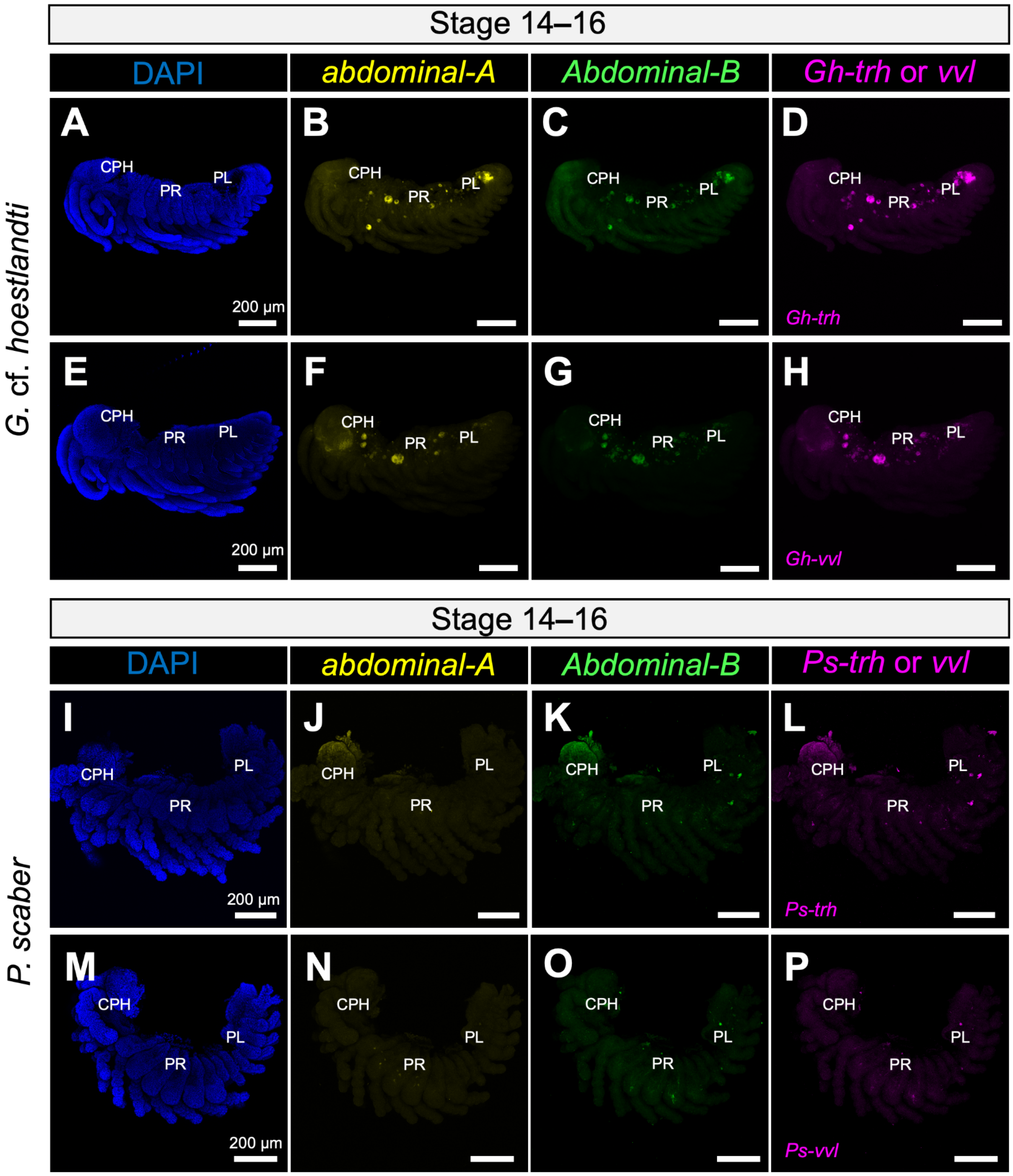


**Figure S7. HCR negative controls**

Negative-control images from the multiplex HCR experiments. The panels show DAPI (A, E, I, M), the detection channel corresponding to *abdA* (B, F, J, N), the channel corresponding to *AbdB* (C, G, K, O), and the channels corresponding to *trh* (D, L) or *vvl* (H, P). Although non-specific fluorescence remains visible in some internal structures (mainly in yolk), no distinct target expression patterns were detected.

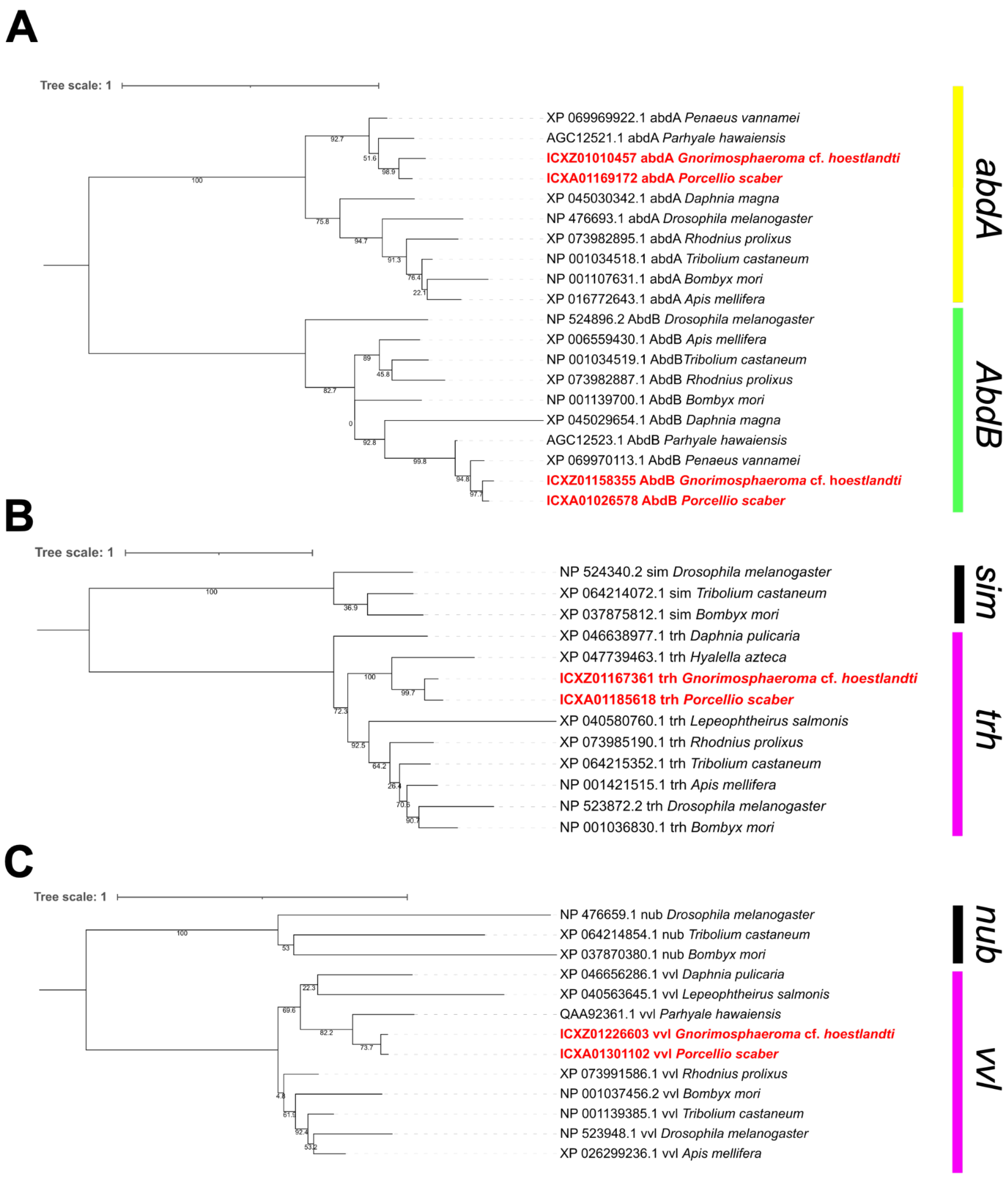


**Figure S8. Phylogenetic placement of candidate isopod sequences**

Gene trees comparing the focal isopod sequences with reference pancrustacean sequences. (A) *abdominal-A* and *Abdominal-B*. (B) *trachealess* and *single-minded*. (C) *ventral veins lacking* and *nubbin*. The focal isopod sequences are shown in red. Numbers at nodes are the support values (ultrafast bootstrap values). Scale bars indicate 1 amino acid substitution per site. Sequence sources are listed in Tables S1 and S2.
